# Changes in urinary proteome in healthy individuals taking multi-vitamin/mineral supplements

**DOI:** 10.1101/2024.06.18.599489

**Authors:** Ziyun Shen, Minhui Yang, Haitong Wang, Yuqing Liu, Youhe Gao

## Abstract

Multivitamin/mineral supplements are widely used in many populations. Urine proteome has not been used to investigate its overall effects on healthy individuals. In this study, the urine proteome changed after taking compound nutrient supplements for 2 weeks and 4 weeks, and the differential proteins and their enriched pathways may be associated with nutrient supplementation. We noted that after taking the supplements for 2 weeks, the amount of erythropoietin receptor in urine was average 449 times higher than before, with four out of nine people experiencing changes from 0 to 14635, 11851, 11181, and 5930. The results of this study revealed that the urine proteome can provide new clues about the relatively short-term health effects of the supplements. It may help optimize guidelines and recommendations for the use of supplements.

## 1 Introduction

Micronutrients and vitamins, collectively known as micronutrients (MNs), are essential for human metabolism. In today’s society, there is a growing concern for maintaining good health. Multivitamin/mineral supplements (MVM) are widely used in many populations^[1]^. Multivitamin/mineral supplements cover a wide range of nutrients, and can provide the body with dietary deficiencies of trace elements and vitamins to meet the body’s multiple nutritional needs. As people become more health-conscious, more and more people are using these supplements with the expectation that they will promote good health by maintaining a well-rounded nutritional state. However, while many people use these supplements to improve their health, much remains unknown about their specific effects on the human body.

Past research has explored the role of multinutrient supplements. In the early years, most of the studies focused on animal experiments, and the population studies mainly focused on the effects of single vitamin or mineral supplementation, and the focus was mostly on disease populations or supplementation treatments for special populations, such as the elderly, pregnant women, and athletes, etc., and there were fewer studies on multinutrient supplementation interventions for the healthy general population. Some studies have shown that multinutrient supplementation can provide micronutrients and vitamins needed to strengthen the immune system, improve cardiovascular health, and increase antioxidant capacity. Multiple studies have shown that there are interactions between vitamins and minerals, which together are involved in a variety of physiological processes, including immune regulation, energy metabolism, and cellular repair. Complex nutrients are important for healthy individuals’ nutritional status, antioxidant capacity^[2]^, exercise capacity, subjective stress and mood^[3]^. The effects of complex nutrients have also been extensively studied. These studies provide some evidence to support the potential benefits of multinutrient supplementation, but often require large population sizes or long experimental times, and we look forward to the urine proteome providing new research perspectives in nutrition that are sensitive to the effects of short-term supplementation on the body. Since urine is not part of the internal environment, in contrast to plasma, there is no mechanism for homeostasis, and it is able to accumulate early changes in the physiological state of the organism, reflecting more sensitively the changes in the organism, and is a source of next-generation biomarkers^[4]^. The proteins in urine contain a wealth of information that can reflect small changes produced in different systems and organs of the organism. Our laboratory has previously reported that the urine proteome can reflect the effects of magnesium threonate intake on the body in a more systematic and comprehensive manner, and has the potential to provide clues for clinical nutrition research and practice^[5]^. To date, no studies have addressed the effects of complex nutrients on the urinary proteome of healthy adults.

The aim of this study was to investigate the changes in the urinary proteome after supplementation with complex nutrients in healthy individuals and to further analyze the potential health effects of such changes. This study will help to reveal the effects of complex nutrient supplementation on human metabolism, physiological functions and potential disease risks, provide new clues for understanding its health effects and developing personalized nutritional intervention strategies in the future, and is expected to provide a new scientific basis for the promotion of the quality of life in healthy populations, as well as provide useful insights into the prevention and management of chronic diseases.

## 2 Materials and Methods

### 2.1 Experimental materials

#### 2.1.1 Experimental consumables

1.5 ml/2 ml centrifuge tube (Axygen, USA), 50 ml/15 ml centrifuge tube (Corning, USA), 96-well cell culture plate (Corning, USA), 10 kD filter (Pall, USA), Oasis HLB solid-phase extraction column (Waters, USA), 1 ml/200 ul/20 ul pipette tips (Axygen, USA), BCA kit (Thermo Fisher Scientific, USA), high pH reverse peptide isolation kit (Thermo Fisher Scientific, USA), iRT (indexed retention time, BioGnosis, UK).

#### 2.1.2 Experimental apparatus

Freezing high-speed centrifuge (Thermo Fisher Scientific, USA), vacuum concentrator (Thermo Fisher Scientific, USA), DK-S22 electric thermostatic water bath (Shanghai Jinghong Experimental Equipment Co., Ltd.), full-wavelength multifunctional enzyme labeling instrument (BMG Labtech, Germany), oscillator (Thermo Fisher Scientific, USA), TS100 constant temperature mixer (Hangzhou Ruicheng Instrument Co., Ltd.), electronic balance (METTLER TOLEDO, Switzerland), −80 □ ultra-low-temperature freezer refrigerator (Thermo Fisher Scientific, U.S.A.), EASY-nLC1200 ultra-high performance liquid Chromatography (Thermo Fisher Scientific, USA), Orbitrap Fusion Lumos Tribird Mass Spectrometer (Thermo Fisher Scientific, USA).

#### 2.1.3 Experimental reagents

The nutrient supplements used were 21 Gold Vita Multi-Vitamin Tablets (Hangzhou Minsheng Health Pharmaceutical Co., Ltd.). In addition, also used Trypsin Golden (Promega, USA), Dithiothreitol DTT (Sigma, Germany), Iodoacetamide IAA (Sigma, Germany), Ammonium Bicarbonate NH4HCO3 (Sigma, Germany), Purified water (Wahaha, China), Methanol for mass spectrometry (Thermo Fisher Scientific, USA), Acetonitrile for mass spectrometry (Thermo Fisher Scientific, USA), Purified water for mass spectrometry (Thermo Fisher Scientific, USA), Tris-Base (Promega, USA) and other reagents.

#### 2.1.4 Analysis software

Proteome Discoverer (Version2.1, Thermo Fisher Scientific, USA), Spectronaut Pulsar (Biognosys, UK), Ingenuity Pathway Analysis (Qiagen, Germany); R studio (Version1.2.5001); Xftp 7; Xshell 7.

### 2.2 Experimental Methods

#### 2.2.1 Study population selection and design

A total of 11 healthy adult volunteers aged 22-27 were selected as experimental subjects, including 5 males and 6 females, and the selection criteria were set as healthy physical examination, excluding heart, lung, liver and spleen disorders. The dietary and lifestyle habits before and after the intervention were kept basically unchanged, and no nutritional supplements or dietary supplements were consumed from one week before the experiment to the beginning of the experiment. Healthy volunteers were given 2 complex nutrient tablets per day for 4 weeks.

The complex nutrient supplement used in the study was 21 Gold Vita Multivitamin tablets. Each healthy volunteer took 2 Complex Nutrient tablets per day for 4 weeks. Each complex nutrient tablet contained L-lysinate 12.5mg, vitamin A 2500IU, vitamin D2 200IU, vitamin E 5mg, vitamin B1 2.5mg, vitamin B2 2.5mg, vitamin B6 0.25mg, vitamin B12 0.5ug, vitamin C 25mg, niacinamide 7.5mg, calcium pantothenate 2.5mg, diazotartaric acid Choline 25mg, Inositol 25mg, Iron 5mg, Iodine 0.05mg, Copper 0.5mg, Manganese 0.5mg, Zinc 0.25mg, Calcium Hydrogen Phosphate 279mg, Magnesium 0.5mg, Potassium 5mg.

**Fig. 1.**
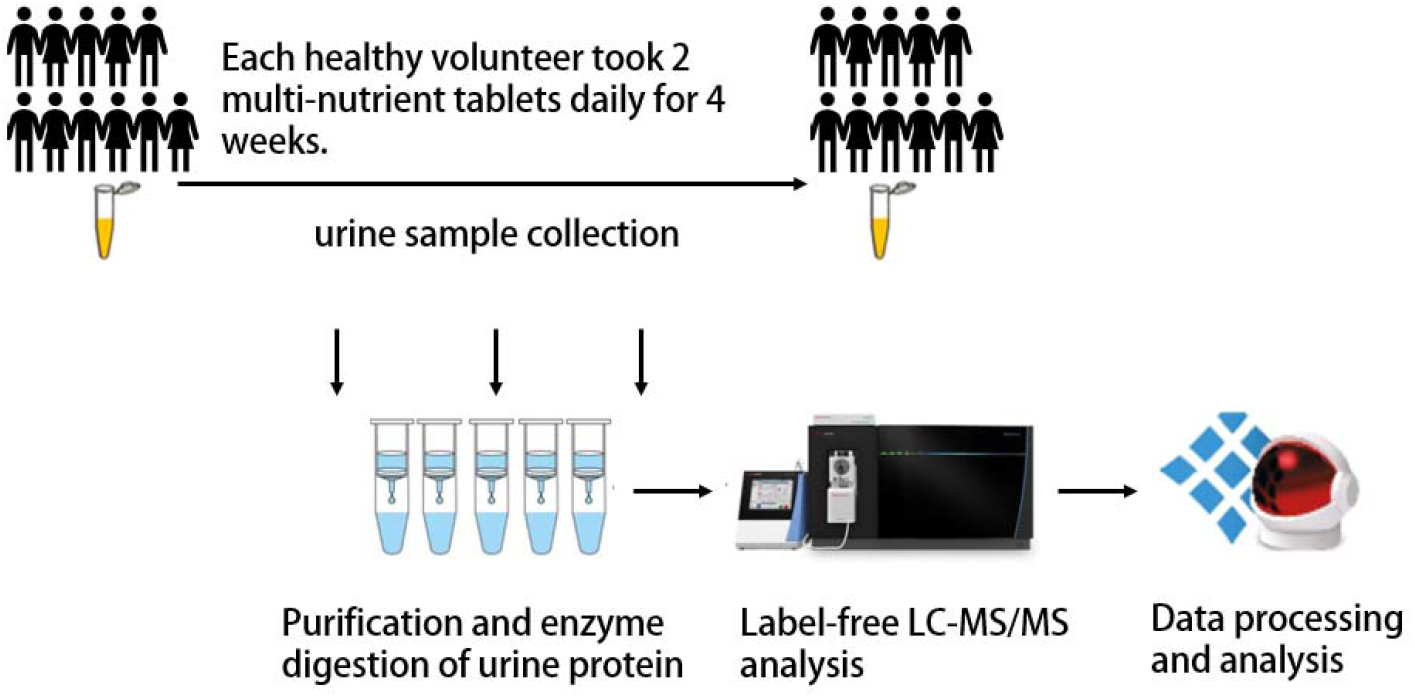
Research methodology and technical route

#### 2.2.2 Urine sample collection

In this study, urine samples were collected from 11 healthy volunteers before taking the multinutrient supplement (labeled as W0, numbered 301-311), after 2 weeks of taking it (labeled as W2, numbered 401-411), and after 4 weeks of taking it (labeled as W4, numbered 501-511). Urine samples were mid-morning urine, avoiding the female menstrual period. Urine samples were collected and placed in a −80°C refrigerator for temporary storage.

#### 2.2.3 Urine sample processing

Remove 2ml of urine samples and thaw, centrifuge at 4◻, 12000×g for 30 minutes, remove cellular debris, take the supernatant and add 1M dithiothreitol (DTT, Sigma) storage solution 40ul, to reach the working concentration of DTT 20mM, mix well and then heated in a metal bath at 37□ for 60 minutes, cool to room temperature, add iodoacetamide (Iodoacetamide, IAA, Sigma) storage solution 100ul, to reach the working concentration of IAM, mix well and then react for 45 minutes at room temperature and protected from light. Iodoacetamide, IAA, Sigma) reservoir solution 100ul, to reach the working concentration of IAM, mix well, and then the reaction was carried out for 45 minutes at room temperature and protected from light. At the end of the reaction, the samples were transferred to new centrifuge tubes, mixed thoroughly with three times the volume of pre-cooled anhydrous ethanol, and placed in a refrigerator at −20°C for 24 h to precipitate the proteins. At the end of precipitation, centrifuge the sample at 4◻ for 30 minutes at 10,000×g, discard the supernatant, dry the protein precipitate, and add 200ul of 20mM Tris solution to the protein precipitate to reconstitute it. After centrifugation, the supernatant was retained and the protein concentration was determined by the Bradford method. Using the filter-assisted sample preparation (FASP) method, urinary protein extracts were added to the filter membrane of a 10kD ultrafiltration tube (Pall, Port Washington, NY, USA), washed three times with 20mM Tris solution, respectively, and the protein was re-solubilized by the addition of 30mM Tris solution, and the protein was added in a proportional manner (urinary protein: trypsin = 50:1) to each sample. Trypsin (Trypsin Gold, Mass Spec Grade, Promega, Fitchburg, WI, USA) was digested and incubated at 37°C for 16 h. The digested filtrate was the peptide mixture. The collected peptide mixture was desalted by an Oasis HLB solid phase extraction column and dried under vacuum, and stored at −80°C. The peptide mixture was then extracted from the peptide mixture with a 0.5μ of 0.1 % PBDE. The lyophilized peptide powder was re-dissolved by adding 30 μL of 0.1% formic acid water, and then the peptide concentration was determined by using the BCA kit, and the peptide concentration was diluted to 0.5 µg/µL, and 4 µL of each sample was taken out as the mix sample.

#### 2.2.4 LC-MS/MS Tandem Mass Spectrometry Analysis

All identification samples were added to a 100-fold dilution of iRT standard solution at a volume ratio of 20:1 sample:iRT, and retention times were standardized. Data-independent acquisition (DIA) was performed on all samples, and each sample was repeated 3 times, with 1-mix samples inserted every 10 pins as quality control. The 1ug samples were separated using EASY-nLC1200 liquid chromatography (elution time: 90min, gradient: mobile phase A: 0.1% formic acid, mobile phase B: 80% acetonitrile), and the eluted peptides were entered into the Orbitrap Fusion Lumos Tribird mass spectrometer for analysis, and the corresponding raw files of the samples were generated.

#### 2.2.5 Data processing and analysis

The raw files collected in DIA mode were imported into Spectronaut software for analysis, and the highly reliable protein standard was peptide q value<0.01. The peak area quantification method was applied to quantify the protein by applying the peak area of all fragmented ion peaks of secondary peptides, and the automatic normalization was processed.

Proteins containing two or more specific peptides were retained, missing values were replaced with 0, the amount of different proteins identified in each sample was calculated, and the samples were compared to screen for differential proteins.

Unsupervised cluster analysis (HCA), principal component analysis (PCA), and OPLS-DA analysis were performed using the Wukong platform (https://omicsolution.org/wkomics/main/). Functional enrichment analysis of differential proteins was performed using the DAVID database (https://david.ncifcrf.gov/) to obtain results in 3 areas: biological process, cellular localization and molecular function. Differential proteins and related pathways were searched based on Pubmed database (https://pubmed.ncbi.nlm.nih.gov/). Protein interaction network analysis was performed using the STRING database (https://cn.string-db.org/).

## 3 Results and Discussion

### 3.1 Results of protein identification of samples

After extraction of urinary proteins, enzymatic digestion, and liquid chromatography tandem mass spectrometry analysis of the collected urine samples, we (counting the mixed samples) identified an average of 3077 proteins (protein and peptide FDR <1%) per sample, with a standard deviation of 628. The average number of proteins identified before taking the multinutrient supplements (notated as W0, numbered 301-311, and 11 samples) was 3026, with a standard deviation of 663. The standard deviation was 663; after 2 weeks of taking the supplement (noted as W2, numbered 401-404, 406-410, with 9 samples), the mean identified protein was 2890, with a standard deviation of 434; after 4 weeks of taking the supplement (noted as W4, numbered 501-511, with 11 samples), the mean identified protein was 3058, with a standard deviation of 657; and the mean identified protein of the MIX samples was 3745 with a standard deviation of 337, indicating good reproducibility of the data.

### 3.2 Changes in urinary proteome after two weeks of nutrient supplement intake

#### 3.2.1 Differential Protein Analysis

Two-tailed, paired t-tests were performed to screen for 228 differential proteins by replacing missing values with 0 and comparing samples before nutrient supplement intake (W0) with samples 2 weeks after nutrient supplement intake (W2). The conditions for screening differential proteins were: p-value <0.05 for t-test analysis, and Fold change (FC) >1.5 or <0.67.

Among them, there were 14 differential proteins with a fold change of >10 and 24 differential proteins with a fold change of >5. Due to space limitation, only the differential proteins with FC >5 or <0.2 are shown in the table.

Erythropoietin receptor has an FC of 449.5. Erythropoietin promotes the proliferation and differentiation of hematopoietic progenitor cells, and erythropoietin and its receptor play neurotrophic and neuroprotective roles in humans^[6]^. Erythropoietin and its receptors play neurotrophic and neuroprotective roles in the human body. Some studies have shown that iron and complex micronutrients significantly increased hemoglobin, erythrocyte pressure volume and serum iron levels in mice. It is possible that supplementation with complex nutrients is associated with erythrocyte^[7]^.

Four differential proteins showed a change from presence to absence with FC of 0, including 26S proteasome non-ATPase regulatory subunit 8, mevalonate diphosphate decarboxylase, procalcitonin γ-A2, and 2-amino-3-carboxymuconate-6-semialdehyde decarboxylase.

#### 3.2.2 Differential Protein Functional Annotation Analysis

Gene Oncology (GO) analysis of 228 differential proteins (p-value <0.05, FC>1.5 or <0.67) using the DAVID database enriched to 74 biological processes (BPs) (p-value <0.05), as shown in the Table 2. Due to space constraints, the top 40 BPs are taken according to p-value size for presentation. 24 molecular functions(MF) are enriched, as shown in the table 3, including protein binding, cadherin binding, GTP binding, GTPase activity, G-protein beta/gamma-subunit complex binding, guanyl nucleotide binding, cytoskeletal protein binding, G-protein coupled receptor binding, GDP binding, ATPase binding, calcium-dependent protein binding, actin binding, ubiquitin binding, protein kinase activator activity, proteoglycan binding, cAMP response element binding, calcium ion binding, transmembrane signaling receptor activity, protein kinase A binding, ATPase activator activity, collagen binding, cell adhesion molecule binding, natriuretic peptide receptor activity, heparin binding.

**Table 1.** Significantly different proteins (FC>5 or <0.2, p-value <0.05) for changes before (W0) and after 2 weeks of nutrient supplement intake (W2)

| Protein | Genes | Protein Descriptions | fc | P |
| --- | --- | --- | --- | --- |

| Accessions |  |  |  |  |
| --- | --- | --- | --- | --- |
| P48556 | PSMD8 | 26S proteasome non-ATPase regulatory subunit 8 | 0 | 0.038649 |
| P53602 | MVD | Diphosphomevalonate decarboxylase | 0 | 0.048419 |
| Q5DRB8 | PCDHGA2 | Protocadherin gamma-A2;Protocadherin gamma-A2 | 0 | 0.031711 |
| Q8TDX5 | ACMSD | 2-amino-3-carboxymuconate-6-semialdehyde decarboxylase | 0 | 0.027933 |
| P22695 | UQCRC2 | Cytochrome b-c1 complex subunit 2, mitochondrial | 0.009509 | 0.049178 |
| O00232 | PSMD12 | 26S proteasome non-ATPase regulatory subunit 12 | 0.030438 | 0.02393 |
| Q13098 | GPS1 | COP9 signalosome complex subunit 1 | 0.033099 | 0.045756 |
| P09110 | ACAA1 | 3-ketoacyl-CoA thiolase, peroxisomal | 0.047178 | 0.033051 |
| Q9Y5F2 | PCDHB11 | Protocadherin beta-11 | 0.050547 | 0.01348 |
| P18850 | ATF6 | Cyclic AMP-dependent transcription factor ATF-6 alpha | 0.054547 | 0.006267 |
| P21217 | FUT3;FUT5 | 3-galactosyl-N-acetylglucosaminide<br>4-alpha-L-fucosyltransferase FUT3 | 0.054779 | 0.034761 |
| Q9Y5X3 | SNX5 | Sorting nexin-5 | 0.072226 | 0.043244 |
| Q9UJA9 | ENPP5 | Ectonucleotide pyrophosphatase/phosphodiesterase family member 5 | 0.095243 | 0.032667 |
| O75019 | LILRA1 | Leukocyte immunoglobulin-like receptor subfamily A member 1 | 0.111365 | 0.030126 |
| Q9BQA1 | WDR77 | Methylosome protein 50;Isoform 2 of Methylosome protein 50 | 0.113783 | 0.046788 |
| Q14353 | GAMT | Guanidinoacetate N-methyltransferase | 0.132321 | 0.006693 |
| O43242 | PSMD3 | 26S proteasome non-ATPase regulatory subunit 3 | 0.13622 | 0.008648 |
| Q14264 | ERV3-1 | Endogenous retrovirus group 3 member 1 Env polyprotein | 0.178283 | 0.033023 |
| Q9Y303 | AMDHD2 | N-acetylglucosamine-6-phosphate deacetylase | 0.185786 | 0.03972 |
| O94813 | SLIT2 | Slit homolog 2 protein | 0.189666 | 0.044398 |
| Q9Y265 | RUVBL1 | RuvB-like 1;Isoform 2 of RuvB-like 1 | 0.189757 | 0.031234 |
| Q13428 | TCOF1 | Treacle protein;Isoform 2 of Treacle protein | 7.310235 | 0.046664 |
| P19235 | EPOR | Erythropoietin receptor | 449.5454 | 0.04668 |

**Table 2.** Biological processes (BPs) enriched to differential proteins between groups before (W0) and 2 weeks after (W2) nutrient supplement intake (BPs ranked in the top 40 according to the size of the p-value are shown)

| Term | Count | % | P-Value |
| --- | --- | --- | --- |
| regulation of cell shape | 12 | 5.6 | 5.50E-07 |
| ubiquitin-dependent protein catabolic process via the multivesicular body sorting pathway | 7 | 3.2 | 7.10E-07 |
| protein transport | 19 | 8.8 | 1.00E-06 |
| Rac protein signal transduction | 6 | 2.8 | 7.90E-06 |
| viral budding via host ESCRT complex | 5 | 2.3 | 7.30E-05 |
| multivesicular body assembly | 5 | 2.3 | 2.90E-04 |
| Adenylate cyclase-modulating G-protein coupled receptor signaling pathway | 6 | 2.8 | 3.70E-04 |
| regulation of actin cytoskeleton organization | 7 | 3.2 | 6.30E-04 |
| membrane fission | 5 | 2.3 | 7.80E-04 |
| cell-matrix adhesion | 7 | 3.2 | 1.00E-03 |
| regulation of organelle assembly | 3 | 1.4 | 1.10E-03 |
| endocytosis | 9 | 4.2 | 1.20E-03 |
| establishment of endothelial barrier | 4 | 1.9 | 1.30E-03 |
| negative regulation of adenylate cyclase activity | 4 | 1.9 | 1.30E-03 |
| regulation of cell size | 4 | 1.9 | 1.90E-03 |
| positive regulation of cell migration | 10 | 4.6 | 2.00E-03 |
| Rho protein signal transduction | 5 | 2.3 | 3.30E-03 |
| positive regulation of early endosome to late endosome transport | 3 | 1.4 | 3.70E-03 |
| viral budding | 3 | 1.4 | 3.70E-03 |
| cell-cell adhesion | 8 | 3.7 | 4.50E-03 |
| positive regulation of protein localization to early endosome | 3 | 1.4 | 4.60E-03 |
| signal transduction | 25 | 11.6 | 5.00E-03 |
| actin filament organization | 7 | 3.2 | 5.80E-03 |
| macroautophagy | 5 | 2.3 | 6.50E-03 |
| cell surface pattern recognition receptor signaling pathway | 3 | 1.4 | 6.70E-03 |
| actin filament bundle assembly | 4 | 1.9 | 6.70E-03 |
| positive regulation of renal sodium excretion | 3 | 1.4 | 7.80E-03 |
| endosomal transport | 5 | 2.3 | 9.00E-03 |
| actin crosslink formation | 3 | 1.4 | 9.10E-03 |
| positive regulation of cellular protein catabolic process | 3 | 1.4 | 9.10E-03 |
| positive regulation of urine volume | 3 | 1.4 | 9.10E-03 |
| negative regulation of cell growth | 6 | 2.8 | 1.00E-02 |
| Protein targeting to vacuole involved in ubiquitin-dependent protein catabolic process via the multivesicular body sorting pathway | 3 | 1.4 | 1.00E-02 |
| G-protein coupled acetylcholine receptor signaling pathway | 3 | 1.4 | 1.00E-02 |
| positive regulation of ruffle assembly | 3 | 1.4 | 1.00E-02 |
| regulation of blood pressure | 5 | 2.3 | 1.10E-02 |
| positive regulation of substrate adhesion-dependent cell spreading | 4 | 1.9 | 1.20E-02 |
| intracellular transport | 4 | 1.9 | 1.60E-02 |
| positive regulation of actin filament polymerization | 4 | 1.9 | 1.60E-02 |
| viral release from host cell | 3 | 1.4 | 1.60E-02 |

**Table 3.**
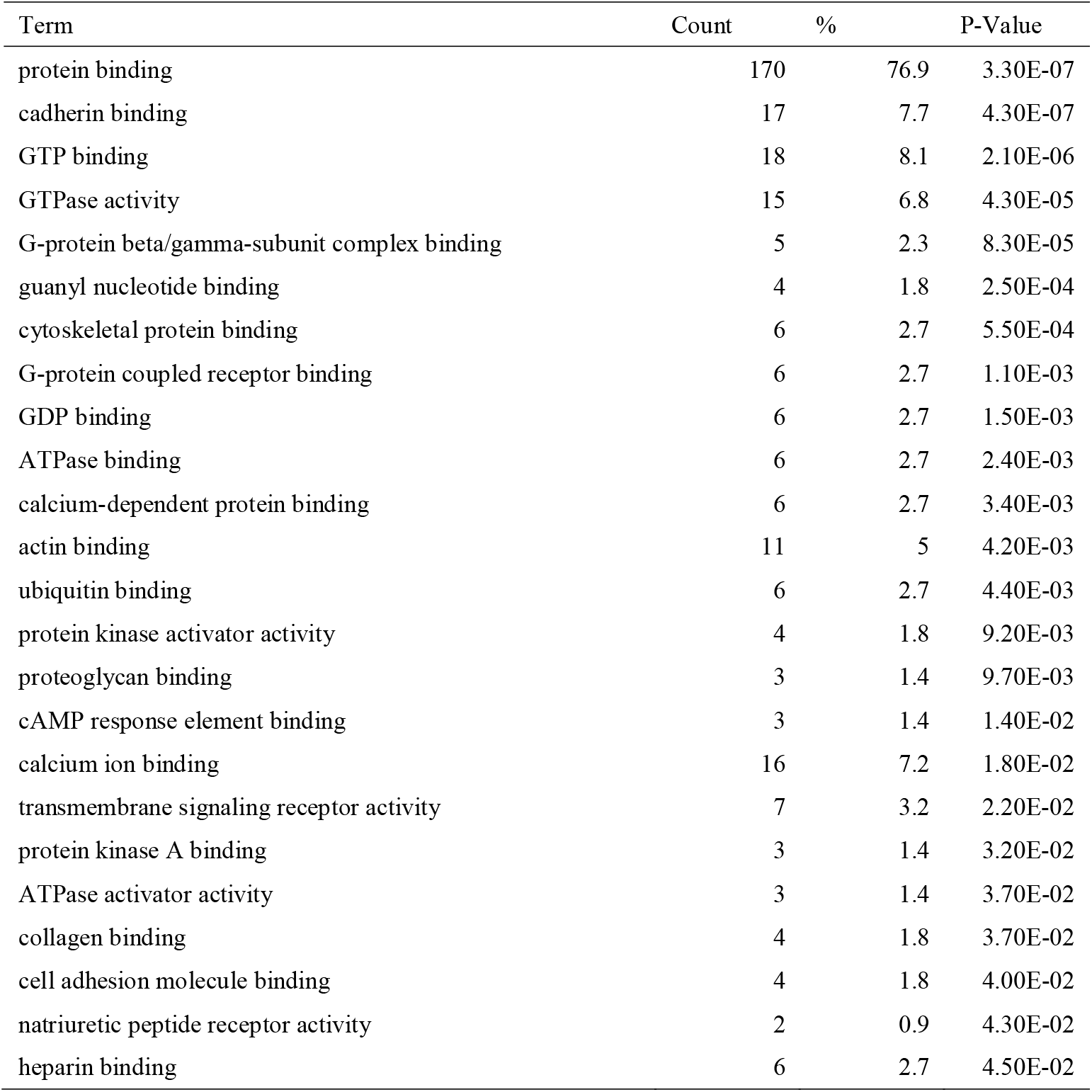
Molecular function(MF) enriched to differential protein between groups before (W0) and after 2 weeks of nutrient supplement intake (W2) (P value <0.05)

| Term | Count | % | P-Value |
| --- | --- | --- | --- |
| protein binding | 170 | 76.9 | 3.30E-07 |
| cadherin binding | 17 | 7.7 | 4.30E-07 |
| GTP binding | 18 | 8.1 | 2.10E-06 |
| GTPase activity | 15 | 6.8 | 4.30E-05 |
| G-protein beta/gamma-subunit complex binding | 5 | 2.3 | 8.30E-05 |
| guanyl nucleotide binding | 4 | 1.8 | 2.50E-04 |
| cytoskeletal protein binding | 6 | 2.7 | 5.50E-04 |
| G-protein coupled receptor binding | 6 | 2.7 | 1.10E-03 |
| GDP binding | 6 | 2.7 | 1.50E-03 |
| ATPase binding | 6 | 2.7 | 2.40E-03 |
| calcium-dependent protein binding | 6 | 2.7 | 3.40E-03 |
| actin binding | 11 | 5 | 4.20E-03 |
| ubiquitin binding | 6 | 2.7 | 4.40E-03 |
| protein kinase activator activity | 4 | 1.8 | 9.20E-03 |
| proteoglycan binding | 3 | 1.4 | 9.70E-03 |
| cAMP response element binding | 3 | 1.4 | 1.40E-02 |
| calcium ion binding | 16 | 7.2 | 1.80E-02 |
| transmembrane signaling receptor activity | 7 | 3.2 | 2.20E-02 |
| protein kinase A binding | 3 | 1.4 | 3.20E-02 |
| ATPase activator activity | 3 | 1.4 | 3.70E-02 |
| collagen binding | 4 | 1.8 | 3.70E-02 |
| cell adhesion molecule binding | 4 | 1.8 | 4.00E-02 |
| natriuretic peptide receptor activity | 2 | 0.9 | 4.30E-02 |
| heparin binding | 6 | 2.7 | 4.50E-02 |

**Table 4.** KEGG pathway enriched by differential proteins between groups before (W0) and 2 weeks after nutrient supplement intake (W2) (P value <0.05)

| Term | Count | % | P-Value |
| --- | --- | --- | --- |
| Endocytosis | 16 | 7.2 | 1.50E-06 |
| Long-term depression | 8 | 3.6 | 1.70E-05 |
| Cholinergic synapse | 10 | 4.5 | 2.40E-05 |
| Regulation of actin cytoskeleton | 13 | 5.9 | 7.00E-05 |
| Growth hormone synthesis, secretion and action | 9 | 4.1 | 2.40E-04 |
| Sphingolipid signaling pathway | 9 | 4.1 | 2.50E-04 |
| Human cytomegalovirus infection | 12 | 5.4 | 2.60E-04 |
| Dopaminergic synapse | 9 | 4.1 | 4.50E-04 |
| Alcoholism | 10 | 4.5 | 1.10E-03 |
| Relaxin signaling pathway | 8 | 3.6 | 2.00E-03 |
| cGMP-PKG signaling pathway | 9 | 4.1 | 2.10E-03 |
| Melanogenesis | 7 | 3.2 | 2.60E-03 |
| Estrogen signaling pathway | 8 | 3.6 | 2.80E-03 |
| Apelin signaling pathway | 8 | 3.6 | 3.00E-03 |
| Parathyroid hormone synthesis, secretion and action | 7 | 3.2 | 3.30E-03 |
| Axon guidance | 9 | 4.1 | 3.60E-03 |
| Parkinson disease | 11 | 5 | 3.60E-03 |
| PI3K-Akt signaling pathway | 13 | 5.9 | 3.80E-03 |
| Prion disease | 11 | 5 | 4.20E-03 |
| Chemokine signaling pathway | 9 | 4.1 | 4.90E-03 |
| Serotonergic synapse | 7 | 3.2 | 5.00E-03 |
| Pathogenic Escherichia coli infection | 9 | 4.1 | 5.90E-03 |
| Gap junction | 6 | 2.7 | 7.20E-03 |
| Human immunodeficiency virus 1 infection | 9 | 4.1 | 8.80E-03 |
| Tight junction | 8 | 3.6 | 8.90E-03 |
| Circadian entrainment | 6 | 2.7 | 1.10E-02 |
| Renin secretion | 5 | 2.3 | 1.50E-02 |
| Retrograde endocannabinoid signaling | 7 | 3.2 | 1.60E-02 |
| Phagosome | 7 | 3.2 | 1.80E-02 |
| Adrenergic signaling in cardiomyocytes | 7 | 3.2 | 1.90E-02 |
| Oxytocin signaling pathway | 7 | 3.2 | 1.90E-02 |
| Cushing syndrome | 7 | 3.2 | 2.00E-02 |
| Leukocyte transendothelial migration | 6 | 2.7 | 2.10E-02 |
| Vasopressin-regulated water reabsorption | 4 | 1.8 | 2.20E-02 |
| MAPK signaling pathway | 10 | 4.5 | 2.30E-02 |
| Rap1 signaling pathway | 8 | 3.6 | 2.60E-02 |
| Platelet activation | 6 | 2.7 | 2.80E-02 |
| Cocaine addiction | 4 | 1.8 | 3.00E-02 |
| Pathways of neurodegeneration - multiple diseases | 13 | 5.9 | 3.00E-02 |
| Chemical carcinogenesis - reactive oxygen species | 8 | 3.6 | 3.40E-02 |
| cAMP signaling pathway | 8 | 3.6 | 3.50E-02 |
| Vascular smooth muscle contraction | 6 | 2.7 | 3.70E-02 |
| Tuberculosis | 7 | 3.2 | 3.80E-02 |
| Human papillomavirus infection | 10 | 4.5 | 3.90E-02 |
| Adherens junction | 5 | 2.3 | 3.90E-02 |
| GnRH signaling pathway | 5 | 2.3 | 3.90E-02 |
| Thermogenesis | 8 | 3.6 | 4.10E-02 |
| Ras signaling pathway | 8 | 3.6 | 4.40E-02 |
| Aldosterone synthesis and secretion | 5 | 2.3 | 4.60E-02 |

**Table 5.**
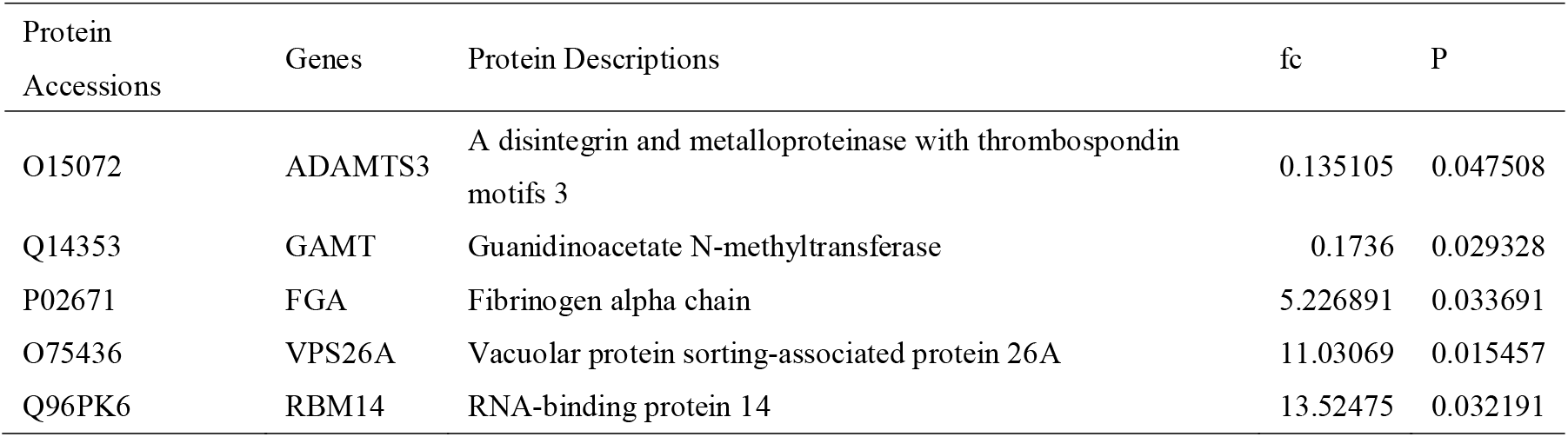
Significantly different proteins (FC>5 or <0.2, p-value <0.05) for changes before (W0) and after 4 weeks of nutrient supplement intake (W4)

| Protein Accessions | Genes | Protein Descriptions | fc | P |
| --- | --- | --- | --- | --- |
| O15072 | ADAMTS3 | A disintegrin and metalloproteinase with thrombospondin motifs 3 | 0.135105 | 0.047508 |
| Q14353 | GAMT | Guanidinoacetate N-methyltransferase | 0.1736 | 0.029328 |
| P02671 | FGA | Fibrinogen alpha chain | 5.226891 | 0.033691 |
| O75436 | VPS26A | Vacuolar protein sorting-associated protein 26A | 11.03069 | 0.015457 |
| Q96PK6 | RBM14 | RNA-binding protein 14 | 13.52475 | 0.032191 |

**Table 6.** Biological processes (BP) enriched to differential proteins between groups before (W0) and after 4 weeks of nutrient supplement intake (W4)

| Term | Count | % | P-Value |
| --- | --- | --- | --- |
| protein deglycosylation | 3 | 3.8 | 3.50E-04 |
| epithelial cell differentiation | 5 | 6.4 | 4.80E-04 |
| protein secretion | 4 | 5.1 | 1.60E-03 |
| intermediate filament organization | 4 | 5.1 | 1.90E-03 |
| epidermis development | 4 | 5.1 | 3.70E-03 |
| intracellular protein transport | 6 | 7.7 | 7.30E-03 |
| negative regulation of extrinsic apoptotic signaling pathway | 3 | 3.8 | 7.80E-03 |
| ER to Golgi vesicle-mediated transport | 4 | 5.1 | 1.30E-02 |
| positive regulation of epidermis development | 2 | 2.6 | 1.80E-02 |
| centriole assembly | 2 | 2.6 | 2.10E-02 |
| endocytic recycling | 3 | 3.8 | 2.20E-02 |
| negative regulation of centriole replication | 2 | 2.6 | 2.80E-02 |
| cell-cell adhesion | 4 | 5.1 | 3.30E-02 |
| endocytosis | 4 | 5.1 | 3.50E-02 |
| mannose metabolic process | 2 | 2.6 | 3.50E-02 |
| protein processing | 3 | 3.8 | 3.80E-02 |
| positive regulation of intracellular signal transduction | 2 | 2.6 | 3.90E-02 |
| retrograde transport, endosome to Golgi | 3 | 3.8 | 4.00E-02 |
| negative regulation of axon regeneration | 2 | 2.6 | 4.20E-02 |
| keratinization | 3 | 3.8 | 4.30E-02 |

**Table 7.** Molecular functions (MFs) enriched for differential proteins between groups before (W0) and after 4 weeks of nutrient supplement intake (W4)

| Term | Count | % | P-Value |
| --- | --- | --- | --- |
| structural constituent of epidermis | 4 | 5.1 | 3.30E-04 |
| GDP binding | 4 | 5.1 | 3.00E-03 |
| cytokine binding | 3 | 3.8 | 1.30E-02 |
| GTP binding | 6 | 7.7 | 1.80E-02 |
| calcium ion binding | 8 | 10.3 | 2.30E-02 |
| protein binding | 56 | 71.8 | 3.30E-02 |
| alpha-mannosidase activity | 2 | 2.6 | 3.30E-02 |
| GTPase activity | 5 | 6.4 | 4.70E-02 |

Differential proteins were enriched to 49 KEGG pathways (P-value < 0.05), including Endocytosis, Long-term depression, Cholinergic synapse, Regulation of actin cytoskeleton, Growth hormone synthesis, secretion and action, Sphingolipid signaling pathway, Human cytomegalovirus infection, Dopaminergic synapse, Alcoholism, Relaxin signaling pathway, cGMP-PKG signaling pathway, Melanogenesis, Estrogen signaling pathway, Apelin signaling pathway, Parathyroid hormone synthesis, secretion and action, Axon guidance, Parkinson disease, PI3K-Akt signaling pathway, Prion disease, Chemokine signaling pathway, Serotonergic synapse, Pathogenic Escherichia coli infection, Gap junction, Human immunodeficiency virus 1 infection, Tight junction, Circadian entrainment, Renin secretion, Retrograde endocannabinoid signaling, Phagosome, Adrenergic signaling in cardiomyocytes, Oxytocin signaling pathway, Cushing syndrome, Leukocyte transendothelial migration, Vasopressin-regulated water reabsorption, MAPK signaling pathway, Rap1 signaling pathway, Platelet activation, Cocaine addiction, Pathways of neurodegeneration - multiple diseases, Chemical carcinogenesis - reactive oxygen species, cAMP signaling pathway, Vascular smooth muscle contraction, Tuberculosis, Human papillomavirus infection, Adherens junction, GnRH signaling pathway, Thermogenesis, Ras signaling pathway, Aldosterone synthesis and secretion.

KEGG pathways such as chronic depression, dopaminergic synapses, synaptic guidance, Parkinson’s disease, serotonergic synapses, and retrograde endogenous cannabinoid signaling show relevance to cognition and mood. It has been shown that nutrients such as Omega-3 fatty acids, antioxidants (vitamin C and zinc), members of the B family of vitamins (vitamin B12 and folate) and magnesium prevent the ways in which oxidative damage to mitochondria and lipids in neuronal circuits associated with cognitive and emotional behaviors^[8]^. 10 differential proteins enriched to cholinergic synapses in this KEGG pathway. Volunteers took a multinutrient supplement containing choline bitartrate (50 mg/d). In addition. some KEGG pathways are related to the immune system.

### 3.3 Changes in urinary proteome after four weeks of ingesting nutrient supplements

#### 3.3.1 Differential protein analysis

Seventy-nine differential proteins were screened by replacing the missing values with 0 and comparing the samples before (W0) and after (W4) 4 weeks of nutrient supplement intake. Screening conditions for differential proteins were: t-test analysis P value < 0.05, Fold change (FC) > 1.5 or < 0.67.

The five proteins with more than a fivefold change in pre-post change were A disintegrin and metalloproteinase with thrombospondin motifs 3, Guanidinoacetate N-methyltransferase, Fibrinogen alpha chain, Vacuolar protein sorting-associated protein 26A, RNA-binding protein 14.

#### 3.3.2 Differential protein function annotation analysis

Gene Oncology (GO) analysis of individual differential proteins (P-value <0.05, FC>1.5 or <0.67) using the DAVID database enriched 20 biological processes (BPs) (P-value <0.05), including protein deglycosylation, epithelial cell differentiation, protein secretion, intermediate filament organization, and epidermal development, intracellular protein transport, negative regulation of exogenous apoptosis signaling pathway, endoplasmic reticulum to Golgi vesicle-mediated transport, positive regulation of epidermal development, centriole assembly, endocytosis recirculation, negative regulation of centriole replication, intercellular adhesion, endocytosis, mannose metabolism process, protein processing, positive regulation of intracellular signaling, retrograde transport, endosome to Golgi, negative regulation of axon regeneration, keratinization.

Differential proteins were enriched to 8 molecular functions (MFs) (P value < 0.05), including structural components of the epidermis, GDP binding, cytokine binding, GTP binding, calcium ion binding, protein binding, α-mannosidase activity, and GTPase activity.

Differential proteins were enriched to 2 KEGG pathways, 6 differential proteins were enriched to endocytosis, and 3 differential proteins were enriched to amino acid biosynthesis.

## 4 Perspective

The results of the study illustrate some changes in the urinary proteome of healthy individuals who continuously took a multinutrient supplement for one month. For example, the average fold change (FC) of Erythropoietin receptor in the urinary proteome after two weeks of taking the complex nutrient was 449.5, with four out of nine people experiencing a change from none (according to the spectronaut’s analysis, it was from 0 to 14635, 11851, 11181, and 5930).

But the individual variability of this change is large. The reasons for this may be due to the following: first, it may be because different people have different dietary habits, different background levels of body nutrient reserves, different types and doses of nutrients need to be supplemented, and the physical condition of different people and the effects of nutrients on the body after intake of multivitamin/mineral supplements are also different, so there is a greater degree of heterogeneity; secondly, it may be that the dosage of multivitamin/mineral supplements is small, and the supplementation time is short and has insignificant impact on the body.

Different individuals may have different nutrient types and dosage needs, it is important to provide personalized supplementation recommendations that are tailored to specific individual, and that personalization can maximize potential benefits and minimize potential risks. The urine proteome also has the potential to provide clues for personalized nutrient supplementation recommendation.

In addition, the changes in the urinary proteome were more significant after two weeks of taking the complex nutrient supplement (W2) than after four weeks of taking the complex nutrient supplement (W4), which may be due to the decrease in compliance of the volunteers with the increase in time, and may also suggest that the nutrients have a more significant effect on the body in the short term, and that the effect of the complex nutrients on the body is not sufficiently pronounced with the increase in time.

The results of this study provide new clues about the health effects of multinutrient supplements from a urinary proteomic perspective, which can help optimize guidelines and recommendations for the use of multinutrient supplements. The personalized analytical methods used in this paper also provide some references for research in related fields. We expect that subsequent studies can measure micronutrients in body fluid samples, various body indicators, and further analyze the changes in the urinary proteome after supplementation with complex nutrients, which can lead to a better understanding of the modulating effects of these supplements on the body’s metabolism, physiological functions, and health status, and provide a basis for the development of new nutritional intervention strategies and innovative functional foods.

